# Net charges of the ribosomal proteins of the *S10* and *spc* clusters of halophiles are inversely related to the degree of halotolerance

**DOI:** 10.1101/2021.11.17.468896

**Authors:** Madhan R Tirumalai, Daniela Anane-Bediakoh, Siddharth Rajesh, George. E. Fox

## Abstract

Net positive charge(s) on ribosomal proteins (r-proteins) have been reported to influence the assembly and folding of ribosomes. A high percentage of r-proteins from extremely halophilic archaea are known to be acidic or even negatively charged. Those proteins that remain positively charged are typically far less so. Herein the analysis is extended to the non-archaeal halophilic bacteria, eukaryotes and halotolerant archaea. The net charges (pH 7.4) of r-proteins that comprise the *S10-spc* operon/cluster from individual microbial and eukaryotic genomes were estimated and intercompared. It was observed that as a general rule, as the salt tolerance of the bacterial strains increased from 5 to 15%, the net charges of the individual proteins remained mostly basic. The most striking exceptions were the extremely halophilic bacterial strains, *Salinibacter ruber* SD01, *Acetohalobium arabaticum* DSM 5501 and *Selenihalanaerobacter shriftii* ATCC BAA-73, which are reported to require a minimum of 18%-21% of salt for their growth. All three strains have a higher number of acidic *S10-spc* cluster r-proteins than what is seen in the moderate halophiles or the halotolerant strains. Of the individual proteins, only uL2 never became acidic. uS14 and uL16 also seldom became acidic. The net negative charges on several of the *S10-spc* cluster r-proteins is a feature generally shared by all extremely halophilic archaea and bacteria. The *S10-spc* cluster r-proteins of halophilic fungi and algae (eukaryotes) were exceptions. They were positively charged despite the halophilicity of the organisms.

**Importance:** The net charges (at pH 7.4) of the ribosomal proteins (r-proteins) that comprise the *S10-spc* cluster show an inverse relationship with the halophilicity/halotolerance levels in both bacteria and archaea. In non-halophilic bacteria, the *S10-spc* cluster r-proteins are generally basic (positively charged), while the rest of the proteomes in these strains are generally acidic. On the other hand, the whole proteomes of the extremely halophilic strains are overall negatively charged including the *S10-spc* cluster r-proteins. Given that the distribution of charged residues in the ribosome exit tunnel influences co-translational folding, the contrasting charges observed in the *S10-spc* cluster r-proteins has potential implications for the rate of passage of these proteins through the ribosomal exit tunnel. Furthermore, the universal protein uL2 which lies in the oldest part of the ribosome is always positively charged irrespective of the strain/organism it belongs to. This has implications for its role in the prebiotic context.

## Introduction

The ribosome is a universal molecular machine comprised of RNA and proteins (1), that catalyzes coded protein synthesis in all three Domains of life (2-4). Thirty-four ribosomal proteins (r-proteins) are universally conserved (5-9). Of these, 21 are encoded by two large clusters, which are analogous to the *S10* and *spc* operons in *E. coli*. These clusters contain four additional genes in *Archaea* and *Eukarya* (9). Given that RNA is negatively charged, the electrostatic properties of r-properties are expected to play a role in stabilizing r-protein-rRNA interactions in the ribosome structure.

In previous examinations of the electrostatic properties of r-proteins, it was observed that extremely halophilic archaeal r-proteins were observed in general to be negatively charged. This is in stark contrast with those from non-halophilic *Archaea* (10, 11). The proteomes of halophilic species are over-represented by acidic residues (12, 13). It is thought that this may reflect a genetic adaptation. Earlier work by Kushner (1978)(14), Kushner and Kamekura (1988)(15), and Ventosa *et al*. (1998)(16), classified halophiles based on their salt requirement and tolerance limit(s). Moderate halophiles have been classified as those growing optimally between 0.5M and 2.5 M salt (14, 15). Strains that are capable of tolerating a broad range of low-high salt concentrations are classified as halotolerant. When growth is possible from low concentration and extends above 2.5M, these strains are classified as extremely halotolerant. Halophiles requiring at least 2M salt for growth, are extreme halophiles (15).

Despite the availability of cultured bacteria, archaea, fungi and algae and their characterized genomes/proteomes (16-32), little is known of the electrostatic properties of the r-proteins of halophilic and halotolerant bacteria, fungi, algae and moderately halophilic archaea. These halophiles range from the moderately halophilic, the halotolerant, the extremely halotolerant, to the extremely obligately halophilic bacteria, fungi and algae (16, 17, 19, 33-67). Additionally, there are several moderately halophilic archaea including several methanogens (20, 39, 68, 69). On the other hand, extremely halophilic strains tend to be obligately halophilic with a minimum salt requirement of 18%-21% salt (70). *A. arabaticum* (18, 71, 72), *S. shriftii* (73) and *S. ruber* (74), with salt requirements of 15%-18% (71), 21% (73) and 20%-25% (75), respectively (Supplementary Table 1), are examples of extremely halophilic bacteria.

Herein, the results of the comparison of the net charges (pH 7.4) of the *S10-spc* r-protein homologs from several halotolerant, extremely halophilic and non-halophilic microbial (bacterial and archaeal) and eukaryotic genomes are reported. The results in part correlate with the extent of halotolerance.

## Methods

Protein and gene sequences from individual microbial (bacterial and archaeal) and eukaryotic organisms were downloaded from the public databases of the National Center for Biotechnology Information (NCBI) (76, 77). The net charges of all proteins at pH 7.4 from each organism were estimated/calculated using the Isoelectric Point Calculator (IPC)(78). The results were cross verified with Prot Pi (https://www.protpi.ch/Calculator/ProteinTool) and PROTEIN CALCULATOR v3.4 (http://protcalc.sourceforge.net/). Additionally, the net charges (at pH 7.4) on the sequences of the r-proteins belonging to the equivalent of the *S10-spc* cluster from each genome were compared with those of the rest of the proteins in each genome.

### Bacterial, archaeal strains and eukaryotes used in comparisons

A list of the organisms (and genomes) covered is given in Supplementary Table 1.

## Results

### Non-halophiles including Archaea and Eukaryotes

The charges on the r-proteins of the homolog equivalents of the *S10-spc* operon/cluster were examined and compared to each other in the dataset of protein sequences from bacteria, archaea, and eukaryotes. In both bacteria and archaea, an increase in the salt tolerance limit is inversely related to the net charges on the r-proteins examined (Fig. 1-4). Based on the general principle that weak acidity is in the pH range 3-6 and strong acidity is <3 (79, 80), the cutoff pH for acidity of the charges was set at 3. As the level of tolerance/halophilicity goes above 15%, many of the r-proteins show charges less than 3.

**Figure 1.**
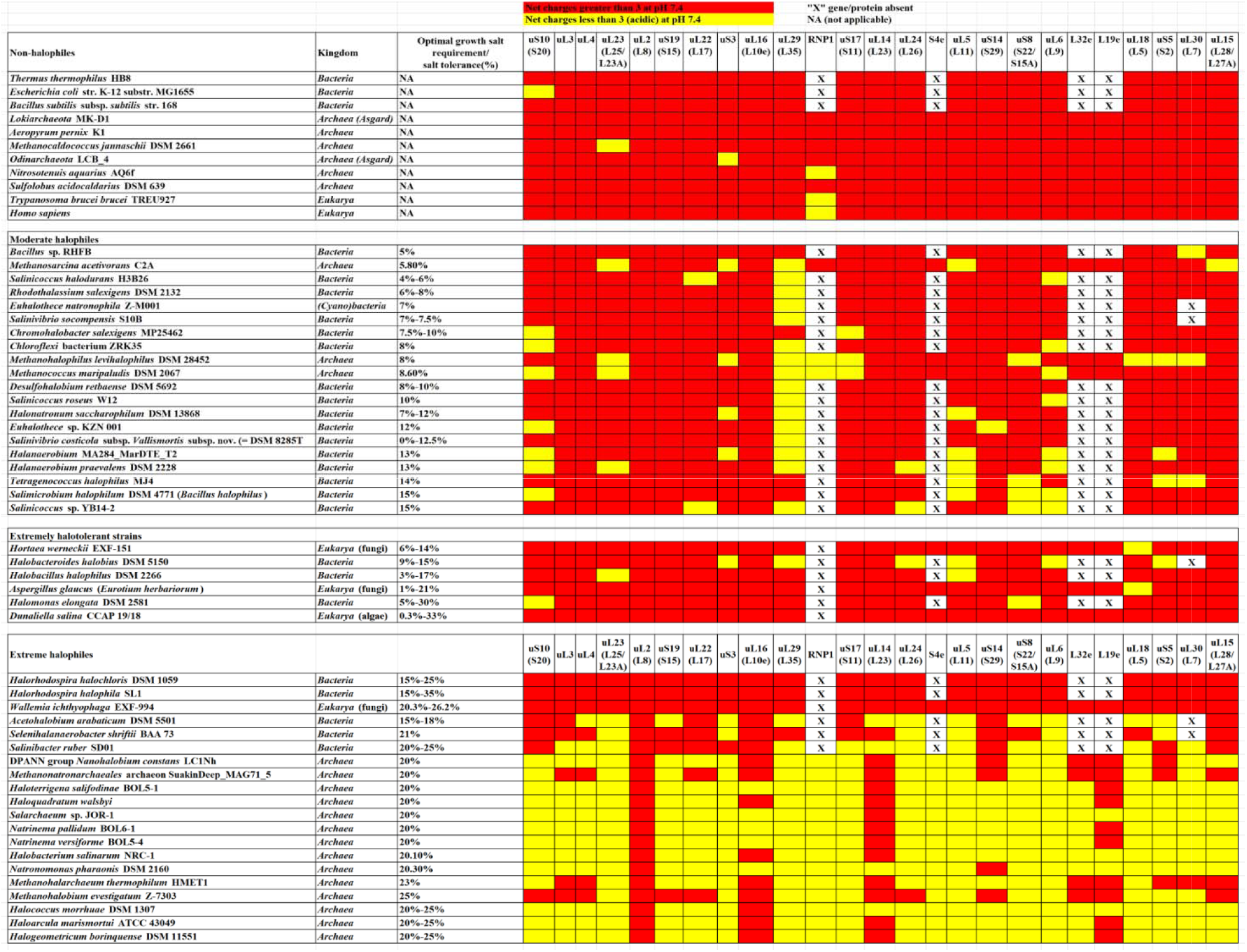
Net charges of the ribosomal proteins of the *S10-spc* cluster from representative strains of *Bacteria, Archaea*, and *Eukarya*. Proteins that have net charges (at pH 7.4) greater than 3 are shown in red 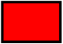, while those with net charges lesser than 3 are in yellow 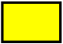.

**Figure 2.**
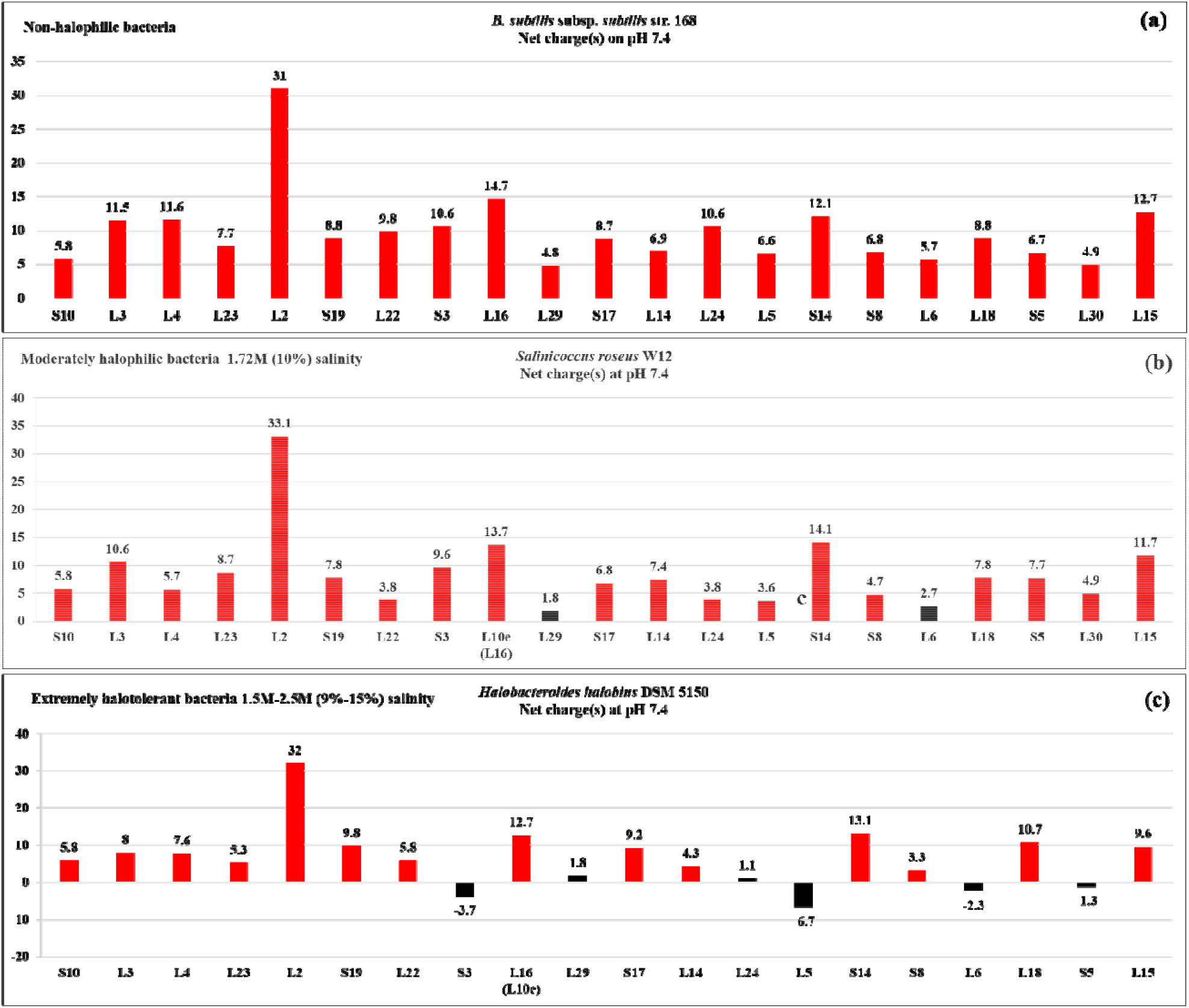
Net charges of the ribosomal proteins of the *S10-spc* cluster from representative strains of moderately halophilic *Bacteria*. (a) *B. subtilis* subsp. *subtilis* str. 168 (non-halophile), (b) *Salinicoccus roseus* W12 (10% salt), and (c) *Halobacteroides halobius* DSM 5150 (9%-15% salt). The charge value of each protein is shown for each bar; charges greater or lesser than three are in red and black respectively.

**Figure 3.**
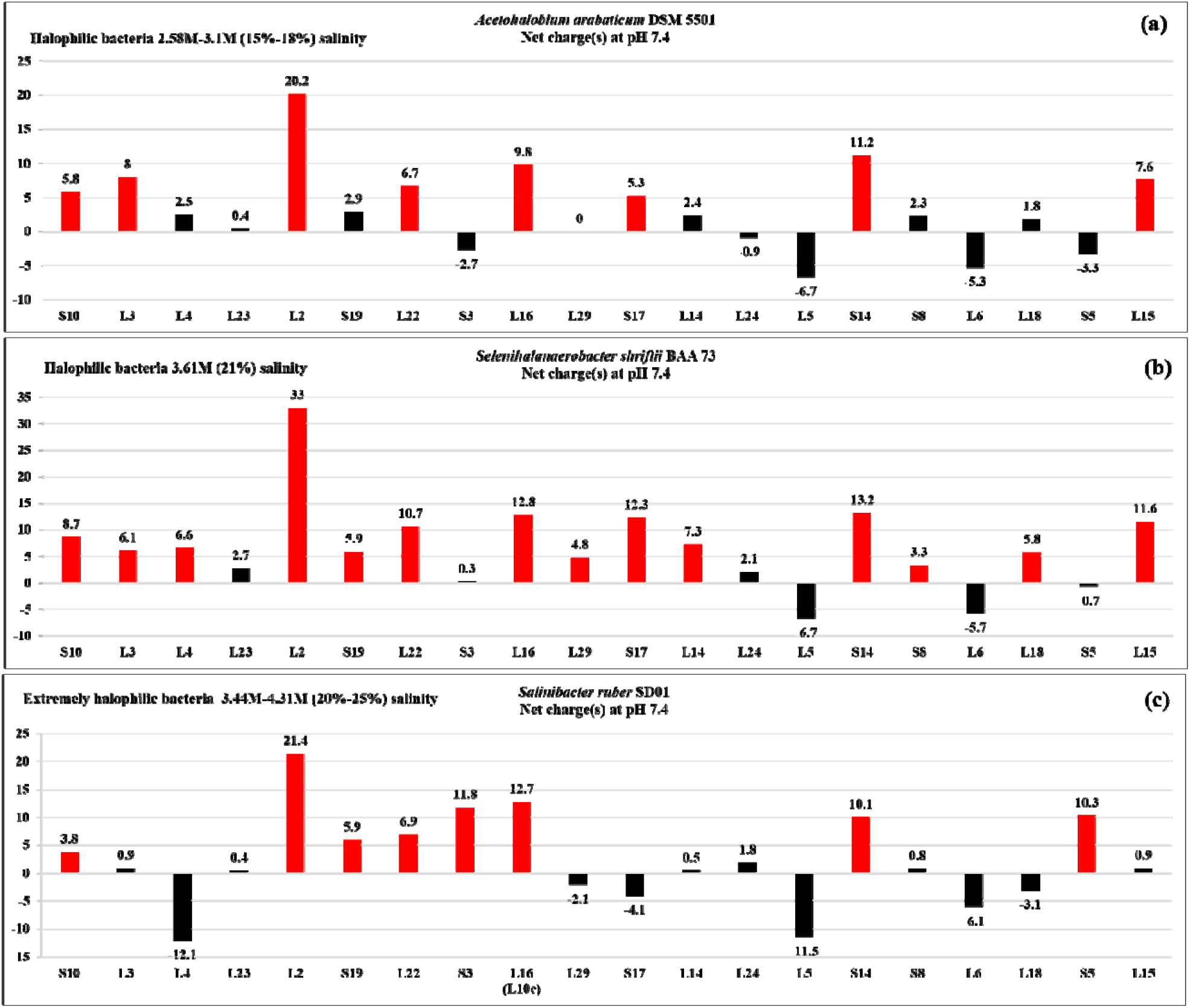
Net charges of the ribosomal proteins of the *S10-spc* cluster from representative strains of extremely halophilic *Bacteria*. (a) *Acetohalobium arabaticum* DSM 5501 (15%-18% salt), (b) *Selenihalanaerobacter shriftii* ATCC BAA-73 (21% salt), and (c) *Salinibacter ruber* SD01 (20%-25% salt). The charge value of each protein is shown for each bar; charges greater or lesser than three are in red and black respectively.

**Figure 4.**
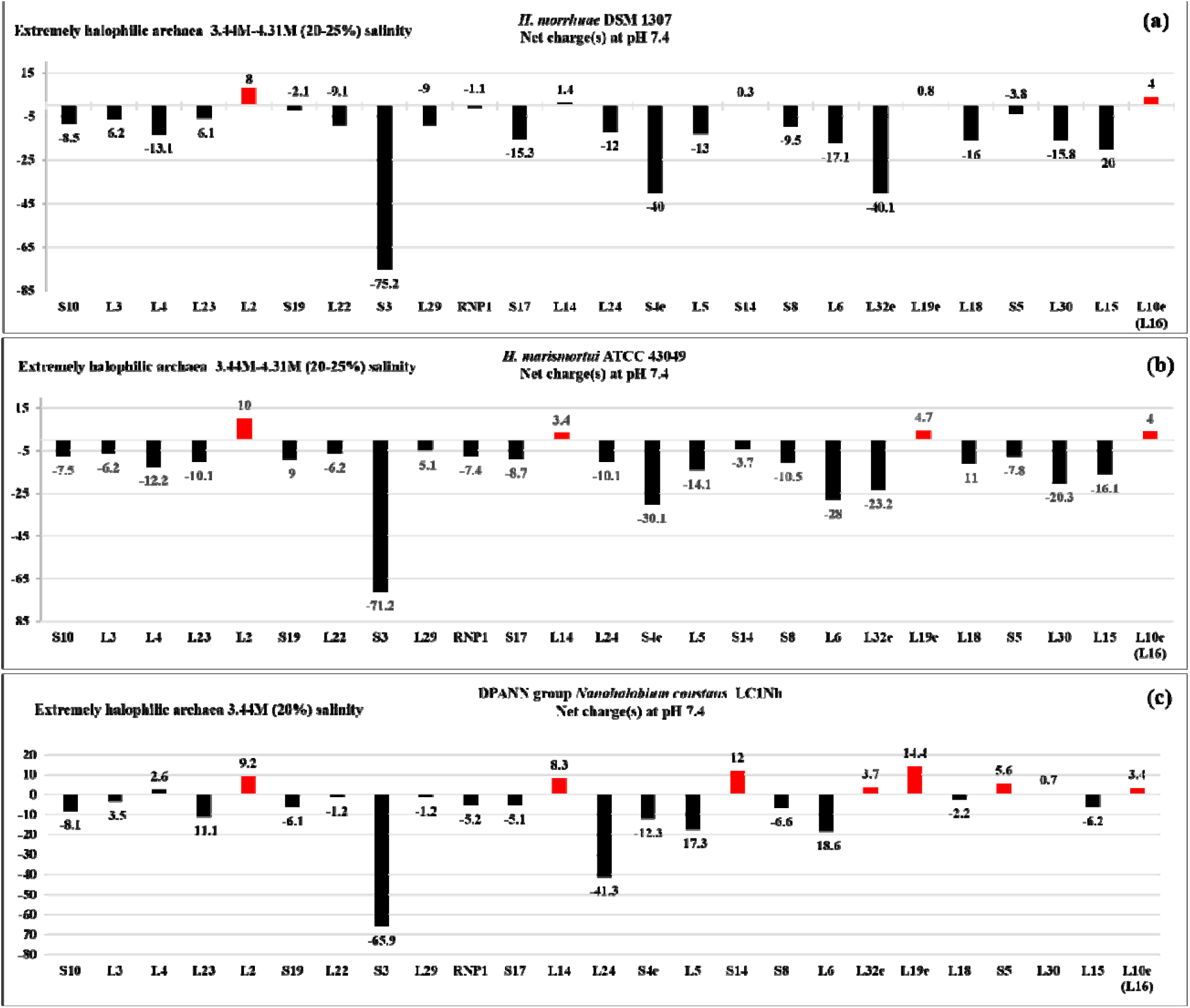
Net charges of the ribosomal proteins of the *S10-spc* cluster from representative strains of extremely halophilic *Archaea*. (a) *H. morrhuae* DSM 1307 (20%-25% salt), (b) *H. marismortui* ATCC 43049 (20%-25% salt), and DPANN group *Nanohalobium constans* LC1Nh (20% salt). The charge value of each protein is shown for each bar; charges greater or lesser than three are in red and black respectively.

### Extremely halophilic archaea

In both the extremely halophilic bacteria and archaea examined, there is a significant increase in the number of *S10-spc* cluster r-proteins that are negatively charged or have charges less than three. Three extremely halophilic bacterial strains *Salinibacter ruber* SD01, *Acetohalobium arabaticum* DSM 5501 and *Selenihalanaerobacter shriftii* BAA-73 were part of this analysis. Twelve of the twenty-one *S10-spc* cluster r-proteins in the strains *S. ruber* SD01 and *A. arabaticum* DSM 5501 and six in *S. shriftii* ATCC BAA-73 possess charges less than 3 (acidic). The acidic properties of the *S10-spc* cluster r-proteins are shared by the halophilic archaea (Fig. 1).

Halotolerant/halophilic fungi (eukarya) and some halotolerant bacteria are exceptions. The bacterial strain *H. elongata* DSM 2581, which is tolerant to a broad range of salt concentrations (5%-30% salt), is extremely halotolerant (81-83). Extremely halophilic bacterial strains *Halorhodospira halochloris* DSM 1059 and *Halorhodospira halophila* SL1 grow optimally at 15%-25% salt and 15%-35% salt, respectively (84-87). In these three bacterial strains, with the exception of uS10 and uS8 in *H. elongata* DSM 2581, all the other *S10-spc* cluster r-proteins have charges greater than 3 (Fig. 1).

The extremely halotolerant fungal strains, namely *Aspergillus glaucus* (*Eurotium herbariorum*), and *Hortaea werneckii* EXF-151, the extremely halophilic fungal strain *W. ichthyophaga* EXF-994, and the extremely halotolerant algae *Dunaliella salina* were all isolated from hypersaline environments (34, 55, 56, 65, 88-94). With the exception of uL18 in *Aspergillus glaucus* and *Hortaea werneckii* EXF-151,the net charges (pH 7.4) of the r-proteins of the *S10-spc* cluster in these organisms are greater than three (Fig. 1 and Supplementary Fig. 1-2). The r-proteins of the *S10-spc* operon cluster in non-halophilic bacteria, non-halophilic archaea and eukarya have positive charges (>3) (Fig. 1). The exceptions were r-proteins uS10 (in the bacterium *E. coli* MG1655), uL23 (in archaeon *M. jannaschii* DSM 2661), uS3 (in the Asgard archaeon *Odinarchaeota* LCB_4), uL16 (L10e) (in the eukaryote *T. brucei brucei* TREU927) and Ribonuclease P1 (RNP1 in the two eukaryotes examined), which had charges less than 3 (Fig.1).

Within the set of r-proteins of the *S10-spc* operon cluster examined, r-protein uL2 was uniquely positively charged (>3) irrespective of the species or the domain to which the species belonged to. In bacteria, uL2 homologs had the highest charges compared to the other proteins of the cluster. In bacteria, the charges on the homologs of uL2 are consistently higher than those of the other proteins in the *S10-spc* cluster. The uL2 homolog with the lowest net charge of 5 (at pH 7.4) is found in the extremely halophilic archaeon *H. salinarum* NRC-1 (Supplementary Table 2). This was also taken into account to set the cutoff value for the net charge(s) of the proteins examined. Likewise, most homologs of uL14 showed charges >3, with the exceptions being the homologs in extreme halophiles *A. arabaticum* DSM 5501 (Bacteria), *S. ruber* SD01 (Bacteria), *N. pharaonis* DSM 2160 (Archaea), and *H. morrhuae* DSM 1307 (Archaea) (Fig. 1-4).

### Net charges on the non-r-proteins (*S10-spc cluster*)

The whole proteomes of the strains that are extremely halophilic are overall negatively charged including the r-proteins. In contrast, an unusual pattern was observed in the proteomes of the non-halotolerant bacteria such as *E. coli*, and *T. thermophilus*, and the moderately halophilic *Bacillus* sp. RHFB (95), *E. natronophila* Z-M001 (cyanobacteria) (41), and *S. roseus* (46). In these strains, while the *S10-spc* cluster proteins show net positive charges, the rest of the proteome showed negative charges (data not shown).

## Discussion

### Salt tolerance and net charges of r-proteins

In a previous analysis of the electrostatic properties of r-proteins from bacteria (*E. coli, T. thermophilus* and *D. radiodurans*), halophilic and non-halophilic archaea, negative charges were uniquely found amongst the r-proteins of the extremely halophilic archaea (10).

In this study, an inverse relationship was observed between the halotolerance limits of bacteria/archaea, and the net charges of the r-proteins of the highly conserved *S10-spc* cluster. In the moderately halophilic bacteria or archaea, which are tolerant up to 15% salt concentration, the charges on the *S10-spc* cluster r-proteins are less than what is found in their homologs in non-halotolerant bacterial strains. The halotolerance is most likely an outcome of properties such as production of intracellular osmolytes, or, solutes, or salting out strategies (70, 89, 96) to counterbalance the ionic imbalance in fluctuating ionic environments. However, in bacteria and archaea, as the halotolerance extends above 15%, many of the *S10-spc* cluster r-proteins of the extremely halophilic bacterial and archaeal strains show net negative charges, implying a genomic level adaptation unique to such strains (Fig. 1-4). In fact, it has been suggested that the halophilic bacterial strain *Salinibacter ruber* SD01 is similar to the extremely halohilic archaeal strains *Halobacterium salinarum* and *Haloarcula marismortui*, both at the genomic and physiological level (97). This gene level adaptation strategy is most likely an evolutionary outcome to minimize energy expenditure that is required to survive in very high salt concentrations.

In contrast to what is observed in the form of drastic changes in the net charges of the r-proteins of the *S10-spc* cluster in extremely halophilic bacteria and archaea, the homologs of the cluster in halophilic fungi and algae (eukaryotes) show positive charges (Fig. 1, Supplementary Fig. 1-2). In an earlier study, it was observed that high contents of acidic residues are frequent in the protein families of three extremely halotolerant/halophilic fungal species, namely, *W. ichthyophaga, H. werneckii* and *E. rubrum* (98). Such strains have also been reported to use traits such as melanin like pigment production, compatible solute production, ion efflux mechanisms, morphological changes and regulation of plasma membrane fluidity to survive in hypersaline conditions (89, 99-102). These observations in halophilic eukaryotes, suggest that the adaptation to hypersaline conditions is likely a result of a combination of acidic residues in proteins as well as changes at the level of physiology and biochemistry. That is clearly not the case with the *S10-spc* cluster protein homologs of the halophilic eukaryotic strains examined in this study.

Of all the *S10-spc* cluster r-proteins, the net charge on uL2 always remains >3. This is despite a steady decrease in the charge(s), corresponding with an increase in the halotolerance, irrespective of the organism/domain (Supplementary Table 2). uL2 is a universal r-protein and is amongst the first set of the large subunit proteins to be assembled into the ribosome (6, 103, 104). In the assembled ribosome structure, it is in very close proximity to what is considered the oldest part of the ribosome, namely, the Peptidyl Transferase Center (PTC) (4, 105, 106). It is also thought to be ancient in origin (107). The universality of the positive charge on all the uL2 homologs has implications for the possible nature of its predecessor peptide in the prebiotic world. In the prebiotic scenario, a positively charged uL2 would have helped maximize stable adhesion/binding to the region surrounding the proto-PTC.

### Implications of net charges on translational rate(s) and ribosome stability

It has been posited that the adaptation to extreme environments such as high salt, might involve structural alterations of proteins, without affecting their functions (108). The ribosome exit tunnel regulates translation and protein folding (109-111). Certain amino acid sequence segments are also known to stall ribosomes (112, 113). In a study on the proteomes of multiple organisms by Requião RD *et al* (114), negatively charged proteins were found to be over-represented. Thus, it was suggested that the charges on the nascent peptide charge is likely one of the factors regulating translation efficiency and protein expression.

However, the vast differences in the net charges of many of the r-proteins of the *S10-spc* cluster between the non-halophiles, moderate and the extreme halophiles, do not appear to significantly affect the ribosome tertiary structure. The core structure of the ribosome is shared by all three domains of life (5).

Furthermore, in studies on halophilic archaea, the stability of ribosomes is known to be severely affected at low-salt concentration buffers (115, 116). However, despite the salt requirement for its stability, the overall basic structure of the ribosome from extreme halophilic archaea such as *H. morrhuae* (116) or *H. marismortui* (117, 118), is similar to those found in bacteria. The amino acid residues are reported to undergo significant intermolecular segregation along the ribosomal proteins, based on their charges. This has been shown to occur in such a manner, so as to have positively and negatively charged residues in buried and solvent export regions, respectively (10). That said, the net low positive or negative charges of halophilic r-proteins found in extremely halophilic archaea/bacteria, could be a major factor contributing to the high salt requirement for the stability of these ribosomes.

Finally, the distribution of charged residues in the ribosome exit tunnel (119) influences the landscape of co-translational folding. For example, the positive charge density of r-proteins in *E. coli* is hypothesized to play a role in the co-translational assembly of ribosomes by delaying the release of nascent r-proteins (120). Electrostatic interactions between the positively charged residues (on the nascent peptide that is being synthesized) and the ribosomal tunnel are known to decrease the translation rate (121). Therefore, it is a moot question as to how negatively charged r-proteins of the *S10-spc* cluster in extreme halophiles affect the rate of the passage of the same through the negatively charged exit tunnel of the ribosome.

## Conclusions

The ribosome in general and the exit tunnel in particular, are primarily negatively charged due to its RNA component which favours positively charged ribosomal proteins. Herein, the charges of the highly conserved ribosomal proteins found in the *S10-spc* gene cluster were used to characterize the extent of halo-tolerance in various microorganisms. It was expected and was found that Bacteria such as *E. coli* or *B. subtilis* that have little to no salt tolerance have the most positively charged proteins. Thus, the charges of the r-proteins of such bacteria are not markedly different from other non-halophilic archaea/eukarya. However, an increase in the salt tolerance limit results in a shift towards a more permanent change in the genome resulting in encoding r-proteins with lower charges. This is evident in the net charges profiles of bacteria capable of growing optimally at salt concentrations above 15%. This trend is similar to what is observed in extremely halophilic archaea. The individual proteins behave differently with uL2 always remaining positive, which may reflect its role in holding the two subunits together. Contrasting charges on the r-proteins in bacteria/archaea, may have implications for the passage of the growing protein through the exit tunnel and thus the translation rate.

## List of abbreviations used

r-protein: ribosomal protein

## Declarations

### 1. Ethics approval and consent to participate

Not Applicable

### 2. Consent for publication

Not applicable

### 3. Availability of data and material

The datasets used and analyzed within the current study are available from the NCBI Website as referenced in the paper.

### 6. Author’s contributions

MRT and GEF conceived and designed the study. MRT, DAB and SR obtained the sequences, estimated the charges on the same and prepared the tables. MRT prepared the figures. MRT, and GEF prepared the manuscript paper which was finalized with help from all the authors. All authors read and approved the final manuscript. SR is a 11^th^ grade student at Clements High school (Class of 2023) who volunteered with the group of Dr. George E. Fox at the University of Houston.

## 7. Acknowledgements

This work was supported by NASA Contract 80NSSC18K1139 under the Center for Origin of Life, Georgia Institute of Technology to GEF.

## Endnotes

None

