## Supplementary Figures 1 and 2 for "Net charges of the ribosomal proteins of the *S10* and *spc* clusters of halophiles are inversely related to the degree of halotolerance"

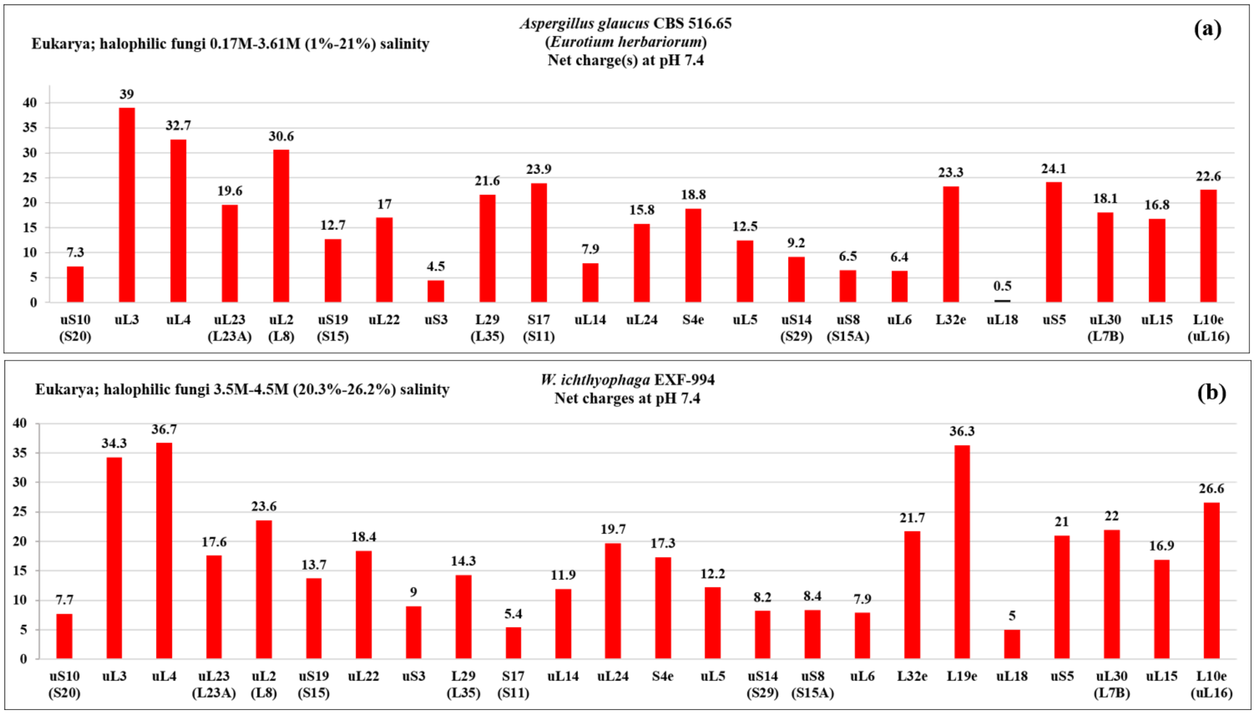


**Supplementary Figure 1.** Net charges of the ribosomal protein homologs of the *S10-spc* cluster from representative strains of halophilic fungi. The charge value of each protein is shown for each bar; charges greater or lesser than three are in red and black respectively (a) *Aspergillus glaucus* CBS 516.65 (*Eurotium herbariorum*) (1%-21% salt), and (b) *W. ichthyophaga* EXF-994 (20.3%-26.2% salt). L19e in *A. glaucus* is truncated (pseudogene) and hence not shown in the figure.

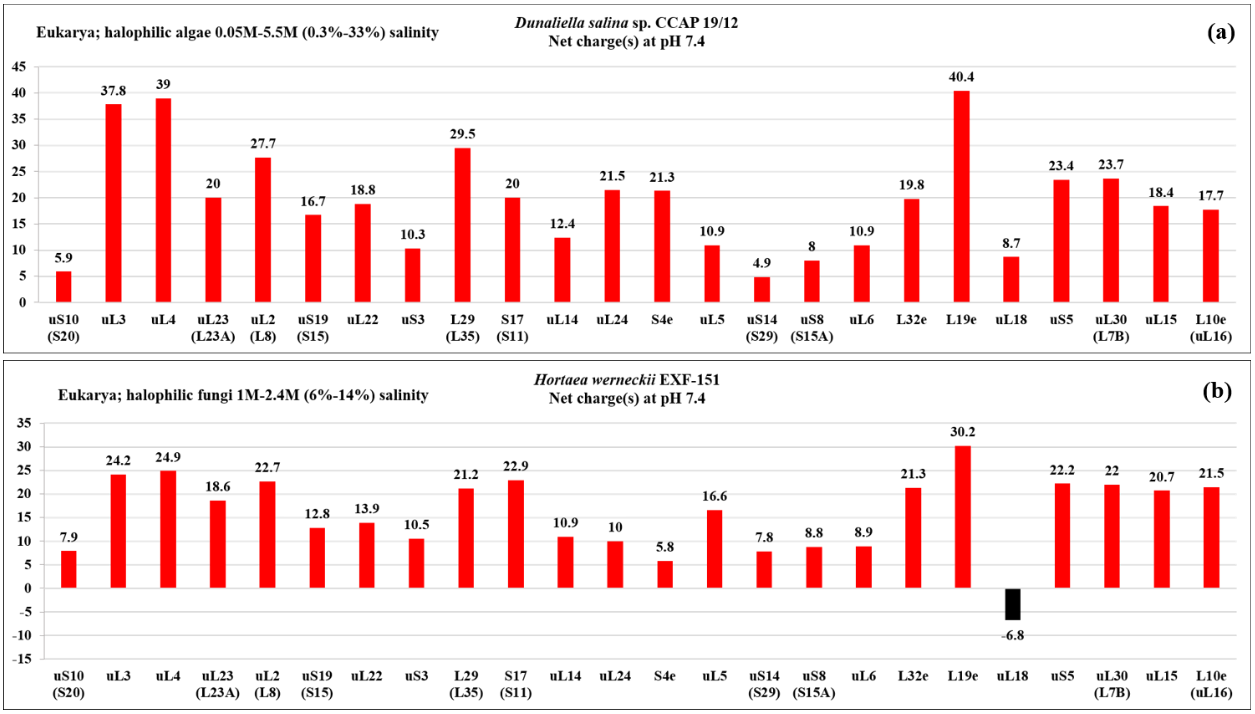


**Supplementary Figure 2.** Net charges of the ribosomal protein homologs of the S10-spc cluster from representative strains of the halophilic (a) algae - *Dunaliella salina* sp. CCAP 19/12 (0.3%-33% salt), and (b) fungi - *Hortaea werneckii* EXF-151 (6%-14% salt). The charge value of each protein is shown for each bar; charges greater or lesser than three are in red and black respectively.
